# Common variants associated with *OSMR* expression contribute to carotid plaque vulnerability, but not to cardiovascular disease in humans

**DOI:** 10.1101/576793

**Authors:** D. van Keulen, I.D. van Koeverden, A. Boltjes, H.M.G. Princen, A.J. van Gool, G.J. de Borst, F.W. Asselbergs, D. Tempel, G. Pasterkamp, S.W. van der Laan

**Affiliations:** Laboratory of Experimental Cardiology, University Medical Center Utrecht, University of Utrecht, Utrecht, The Netherlands; Central Diagnostics Laboratory, University Medical Center Utrecht, University of Utrecht, Utrecht, The Netherlands; Quorics B.V., Rotterdam, The Netherlands; TNO-Metabolic Health Research, Gaubius Laboratory, Leiden, The Netherlands; Translational Metabolic Laboratory, Radboudumc, Nijmegen, The Netherlands; TNO- Microbiology & Systems Biology, Zeist, The Netherlands; Department of Vascular Surgery, University Medical Center Utrecht, University of Utrecht, Utrecht, The Netherlands; Department of Cardiology, Division Heart & Lungs, University Medical Center Utrecht, Utrecht University, Utrecht, The Netherlands; Institute of Cardiovascular Science, Faculty of Population Health Sciences, University College London, London,United Kingdom; Health Data Research UK and Institute of Health Informatics, University College London, London, United Kingdom; SkylineDx B.V., Rotterdam, The Netherlands

**Keywords:** Cardiovascular disease, Atherosclerosis, Plaque, Genetics, OSM, OSMR, LIFR

## Abstract

**Background and aims:** Oncostatin M (OSM) signaling is implicated in atherosclerosis, however the mechanism remains unclear. We investigated the impact of common genetic variants in OSM and its receptors, *OSMR* and *LIFR*, on overall plaque vulnerability, plaque phenotype, intraplaque *OSMR* and *LIFR* expression, coronary artery calcification burden and cardiovascular disease susceptibility.

**Methods and results:** We queried Genotype-Tissue Expression data and found that rs13168867 (C allele) was associated with decreased *OSMR* expression and that rs10491509 (A allele) was associated with increased *LIFR* expression in arterial tissues. No variant was significantly associated with *OSM* expression.

We associated these two variants with plaque characteristics from 1,443 genotyped carotid endarterectomy patients in the Athero-Express Biobank Study. After correction for multiple testing, rs13168867 was significantly associated with an increased overall plaque vulnerability (β=0.118 ± s.e.=0.040, *p*=3.00×10^-3^, C allele). Looking at individual plaque characteristics, rs13168867 showed strongest associations with intraplaque fat (β=0.248 ± s.e.=0.088, p=4.66 × 10^-3^, C allele) and collagen content (β=-0.259 ± s.e.=0.095, p=6.22 × 10^-3^, C allele), but these associations were not significant after correction for multiple testing. rs13168867 was not associated with intraplaque *OSMR* expression. Neither was intraplaque *OSMR* expression associated with plaque vulnerability and no known *OSMR* eQTLs were associated with coronary artery calcification burden, or cardiovascular disease susceptibility. No associations were found for rs10491509 in the *LIFR* locus.

**Conclusions:** Our study suggests that rs1316887 in the OSMR locus is associated with increased plaque vulnerability, but not with coronary calcification or cardiovascular disease risk. It remains unclear through which precise biological mechanisms OSM signaling exerts its effects on plaque morphology. However, the OSM-OSMR/LIFR pathway is unlikely to be causally involved in lifetime cardiovascular disease susceptibility.

## 1. Introduction

The prevalence of cardiovascular disease (CVD) is high, poses a significant global burden and is expected to rise(1). Arterial inflammation, leading to asymmetric focal arterial thickening and atherosclerotic plaque formation and progression, is the primary mechanism underlying CVD(2). Inflammatory cytokines contribute to arterial inflammation and subsequent atherosclerotic plaque formation(3). One cytokine, for which there is mounting evidence suggesting a role in atherosclerosis development is OSM(4,5). It has been shown that OSM is present in both murine and human atherosclerotic plaques(6). Moreover, murine studies showed that OSM receptor (OSMR)^-/-^ApoE^-/-^mice have reduced plaque size and improved plaque stability compared to their *OSMR*-expressing littermates(7), indicating that OSM drives atherosclerosis development. These observations are in line with our previous work, in which we showed that simultaneous signaling of OSM through OSMR and leukemia inhibitory factor receptor (LIFR), induces activation in human endothelial cells, suggestive of a role in atherosclerosis development(8). In contrast, chronic OSM administration to APOE*3Leiden.CETP mice reduces the atherosclerotic lesion size and severity, and high circulating OSM levels correlate with increased post-incident coronary heart disease survival probability in humans(9).

Although all these studies implicate that OSM is involved in atherosclerosis, little is known about the effects of OSM on plaque composition in humans. Grouped in the interleukin 6 subfamily of cytokines, OSM is released by activated immune cells(10–12), and exerts pleiotropic effects on cell proliferation, inflammation, hematopoiesis, tissue remodeling, and development(13). Its signals are transduced through binding to either OSMR or LIFR, which form a heterodimer with glycoprotein 130(8,14), that in turn activates multiple pathways(14). It is suggested that the ratio of the two receptor types expressed on the cell membrane is a potential regulatory mechanism for the multiple, and sometimes opposing, effects that are exerted by OSM(15).

Thus, given its pleiotropic function, it is difficult to predict how OSM contributes to atherosclerotic plaque formation. Cell and murine studies have shown that OSM promotes angiogenesis(4), endothelial activation(8), vessel permeability(16), and osteoblastic differentiation(17). Therefore, increased OSM levels hypothetically results in a higher intraplaque microvessel density, intraplaque hemorrhages and plaque calcification, thereby contributing to plaque destabilization(18,19). In other cell and murine studies, OSM promotes fibroblast proliferation(20), collagen formation(20), smooth muscle cell proliferation(6), and M2 macrophage polarization(21). These processes hypothetically lead to enhanced fibrosis, and attenuated inflammation, thereby contributing to plaque stabilization(22).

Large-scale studies have shown that *cis*-acting genetic variants associated to gene expression (expression quantitative trait loci (eQTLs)(23)) are key to disease susceptibility(24). This means that gene expression in a given tissue differs between individuals carrying different genotypes which ultimately results in differential disease susceptibility. Thus, on the premise that alleles are randomly distributed at conception and are invariant throughout a lifetime, meaning that genetics is not influenced by disease or risk factors, eQTLs can be used as proxies of gene expression to examine the effect on plaque morphology(25). We hypothesized that if circulating *OSM*, or arterial *OSMR* or *LIFR* expression has an effect on plaque morphology, these phenotypic differences will be observed among genotype groups of the eQTL. We aimed to investigate the double-edged sword of OSM signaling on the composition of human atherosclerotic plaques using known eQTLs of circulating *OSM*, and arterial *OSMR* and *LIFR*.

## 2. Materials and methods

### 2.1 Sample collection

The Athero-Express Biobank Study (https://www.atheroexpress.nl) is an ongoing prospective study, containing biomaterial of patients elected for endarterectomy at two Dutch tertiary referral centers. Details of the study design were described before(26). Briefly, blood subfractions are obtained before and arterial plaque material during endarterectomy. Each plaque is dissected into segments of 0.5cm. The culprit lesion is reserved for histological assessment (see below), while surrounding segments are immediately snap frozen in liquid nitrogen and stored at -80°C for later use, e.g., in order to perform RNA-seq (see below). Only carotid endarterectomy (CEA) patients were included in the present study. All research was conducted according to the principles of the Declaration of Helsinki and its later amendments, all patients provided informed consent and the study was approved by the medical ethics committees.

### 2.2 Athero-Express genotyping, quality control, and imputation

Details of genotyping were previously described(26). Briefly, DNA was extracted from EDTA blood or (when no blood was available) plaque samples of 1,858 consecutive patients from the Athero-Express Biobank Study and genotyped in 2 batches. For the Athero-Express Genomics Study 1 (AEGS1), 836 patients, included between 2002 and 2007, were genotyped using the Affymetrix Genome-Wide Human SNP Array 5.0 (SNP5) chip (Affymetrix Inc., Santa Clara, CA, USA). For the Athero-Express Genomics Study 2 (AEGS2), 1,022 patients, included between 2002 and 2013 and not overlapping AEGS1, were genotyped using the Affymetrix Axiom® GW CEU 1 Array (AxM).

Both studies were carried out according to OECD standards. After genotype calling, we adhered to community standard quality control and assurance (QA/QA) procedures of the genotype data from AEGS1 and AEGS2. Samples with low average genotype calling and sex discrepancies (compared to the clinical data available) were excluded. The data was further filtered on 1) individual (sample) call rate >97%, 2) SNP call rate >97%, 3) minor allele frequencies (MAF) >3%, 4) average heterozygosity rate ± 3.0 s.d., 5) relatedness (pi-hat >0.20), 6) Hardy–Weinberg Equilibrium (HWE *p*< 1.0×10^-6^), and 7) population stratification (based on HapMap 2, release 22, b36) by excluding samples deviating more than 6 standard deviations from the average in 5 iterations during principal component analysis (PCA) and by visual inspection as previously described(26). After QA/QA, 657 samples and 403,789 SNPs in AEGS1, and 869 samples and 535,983 SNPs in AEGS2 remained. To correct for genetic ancestry and population stratification we performed PCA in each cleaned dataset to obtain principal components for downstream analyses as described before(26).

We used SHAPEIT2(27) for phasing and finally the data was imputed with 1000G phase 3(28) and GoNL 5(29) as a reference on genome build 37. Note that we only selected the CEA patients in these datasets, leaving 1,443 samples for our further analyses.

### 2.3 Variant selection

We queried data from the Genotype-Tissue Expression (GTEx) Portal (https://gtexportal.org)(23) for *cis*-acting variants (defined as variants within 1Mb of a given gene(30)) that alter *OSM* expression in whole blood, and *OSMR* or *LIFR* expression in non-diseased arterial tissue. We selected common variants with a MAF >3%, which yielded 2 variants in total: rs13168867 for *OSMR* in tibial arterial tissue and rs10491509 for *LIFR* in aortic arterial tissue. We found no eQTL for circulating *OSM* expression, i.e. in whole blood. We harmonized the effect alleles and effect sizes from these eQTLs to match the allele orientation in the Athero-Express Biobank Study data.

### 2.4 Plaque phenotyping

The (immuno)histochemical analysis of plaques have been described previously(26,31,32). Briefly, per plaque, the culprit lesion was identified directly after dissection, fixed in 4% formaldehyde, embedded in paraffin and cut in 5µm sections on a microtome for (immuno)histochemical analysis by pathology experts. Calcification (hematoxylin & eosin, H&E) and collagen content (picrosirius red) were semi-quantitatively scored and defined as no/minor or moderate/heavy. Atheroma size (H&E and picrosirius red) was defined as <10% or ≥10% fat content. Macrophages (CD68) and smooth muscle cells (ACTA2) were quantitatively scored and classified as percentage of plaque area. Intraplaque hemorrhage (H&E) was defined as absent or present, and vessel density was classified as the number of intraplaque vessels (CD34) per 3-4 hotspots.

### 2.5 Plaque vulnerability

Assessment of overall plaque vulnerability was performed as previously described by Verhoeven *et al*(25). Briefly, macrophages and smooth muscle cells were semi-quantitatively defined as no/minor or moderate/heavy. Each plaque characteristic that defines a stable plaque (i.e., no/minor macrophages, moderate/heavy collagen, moderate/heavy smooth muscle cells and <10% fat) was given a score of 0, while each plaque characteristic that defines a vulnerable plaque (i.e., moderate/heavy macrophages, no/minor collagen, no/minor smooth muscle cells and ≥10% fat) was given a score of 1. The score of each plaque characteristic was summed resulting in a final plaque score ranging from 0 (most stable plaque) to 4 (most vulnerable plaque). Intraobserver and interobserver variability were examined previously and showed good concordance (κ=0.6-0.9)(33).

### 2.6 Plaque expression

Detailed information on the RNA sequencing (RNAseq) experiment is described in the Supplemental Material. In short, to assess the global expression profile, plaque segments were thawed, cut up, and homogenized using ceramic beads and tissue homogenizer (Precellys, Bertin instruments, Montigny-le-Bretonneux, France), in the presence of TriPure (Sigma Aldrich), and RNA was isolated according to TriPure manufacturer’s protocol.

Library preparation was performed, adapting the CEL-Seq2 protocol for library preparation(34,35), as described before(36). The primer used for initial reverse-transcription reaction was designed as follows: an anchored polyT, a unique 6bp barcode, a unique molecular identifier (UMI) of 6bp, the 5’ Illumina adapter and a T7 promoter, as described(36). Complementary DNA (cDNA) was then used in the *in vitro* transcription (IVT) reaction (AM1334; Thermo-Fisher). Amplified RNA (aRNA) was fragmented, and cleaned, and RNA yield and quality in the suspension were checked by Bioanalyzer (Agilent). cDNA library construction was initiated according to the manufacturer’s protocol, adding randomhexRT primer as random primer. PCR amplification was done with Phusion High-Fidelity PCR Master Mix with HF buffer (NEB, MA, USA) and a unique indexed RNA PCR primer (Illumina) per reaction. Library cDNA yield and quality were checked by Qubit fluorometric quantification (Thermo-Fisher) and Bioanalyzer (Agilent), respectively. Libraries were sequenced on the Illumina Nextseq500 platform; paired end, 2 × 75bp.

Upon sequencing, retrieved fastq files were de-barcoded, split into forward and reverse reads. Subsequently, these were mapped making use of Burrows-Wheel aligner (BWA(37)) version 0.7.17-r1188 and a cDNA reference (assembly hg19, Ensembl release 84). Read counts and UMI counts were derived from SAM files using custom perl code, and then gathered into count matrices. Genes were annotated with Ensembl ID’s, and basic quality control was performed, encompassing filtering out samples with low gene numbers (<10,000 genes), and read numbers (<18,000 reads). These steps resulted in 641 samples with up to 60,674 genes (Ensembl ID’s), and median of 178,626 reads per sample.

#### Data analysis

Plaque vulnerability scores, and genotypes for rs10491509 and rs13168867, were added to metadata, upon which this was combined with counts and annotation in a SummarizedExperiment object(38). Counts were normalized and transformed making use of the variance stabilization transformation function (vst()) in DESeq2(39). This results in transformed data on a log_2_-scale, normalized for library size, for visualization and ordination purposes. Differential expression analysis between plaque vulnerability scores or genotypes, used as ‘condition variables’ was performed using the DESeq2-function DESeq() on the raw counts. In short, three steps are performed: 1. estimation of size factors, controlling for sequencing depth; 2. estimation of dispersion values, that capture variation around expected values. These expected values take into account sequencing depth and differences caused by variables in the design formula argument, *i*.*e*., ‘design = ∼ condition’ where condition is a variable that specifies which group samples belong to; and 3. fitting a generalized linear model using the above-mentioned size factors and dispersion values, estimating log fold changes. This results in a results table, showing estimated log2 fold changes and p values comparing between two levels of the condition variable. Complete details for statistical procedures used by the DESeq function are described elsewhere(39).

### 2.7 Genetic analyses

Quantitatively scored characteristics (macrophages, smooth muscle cells, and the vessel density) were Box-Cox transformed(40) to obtain a normal distribution. For genetic analyses we used GWASToolKit (https://swvanderlaan.github.io/GWASToolKit/) which is a wrap-around collection of scripts for SNPTEST(41). Continuous and categorical variables were tested using linear and logistic regression models, respectively. Models for genetic analyses were corrected for age, sex, genotyping chip, and genetic ancestry using principal components 1 through 4. Thus, the models were of the form

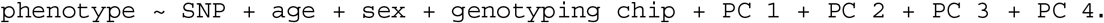

### 2.8 Multiple testing and power

Correction for multiple testing resulted in a corrected p-value of *p*=0.05/((7 plaque phenotypes + plaque vulnerability) × 2 common variants)=3.13×10^-3^. The power of the study was estimated at ±75% based on a sample size of 1,443, a minor allele frequency (MAF) of 0.409 and a relative risk of 1.28 (http://csg.sph.umich.edu/abecasis/cats/gas_power_calculator/).

### 2.9 Data and scripts

Data is available upon request. Scripts are posted at GitHub https://github.com/swvanderlaan/2019_vankeulen_d_osmr.

## 3. Results

### 3.1 Common variants altering OSM, OSMR and LIFR expression

We included and genotyped 1,443 carotid endarterectomy patients in this study. We combined these groups (**Table 1**) for overall plaque vulnerability and phenotype analyses, as we previously showed that the baseline characteristics between the two genotyping groups (AEGS1 and AEGS2) are comparable(26).

**Table 1:**
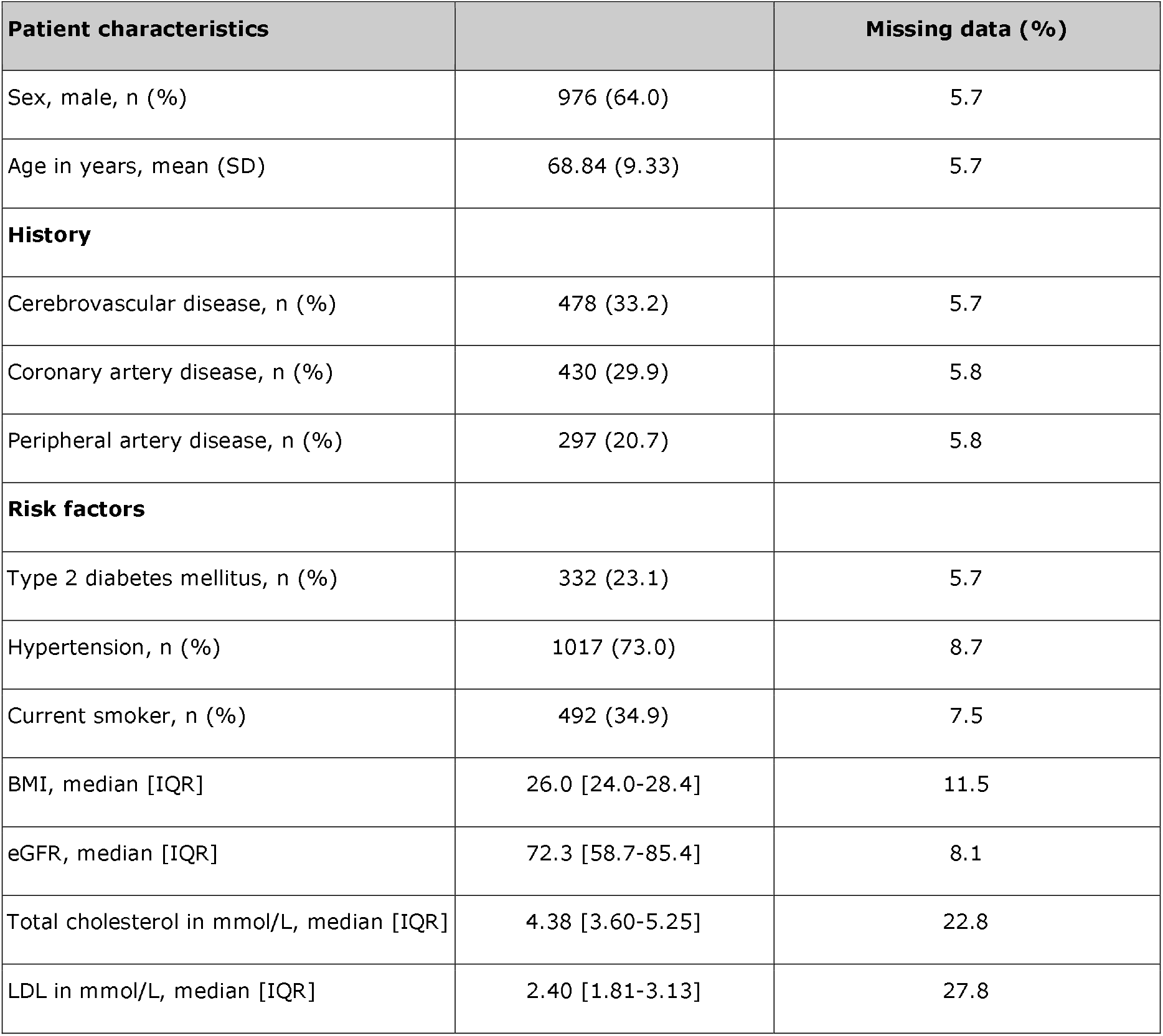
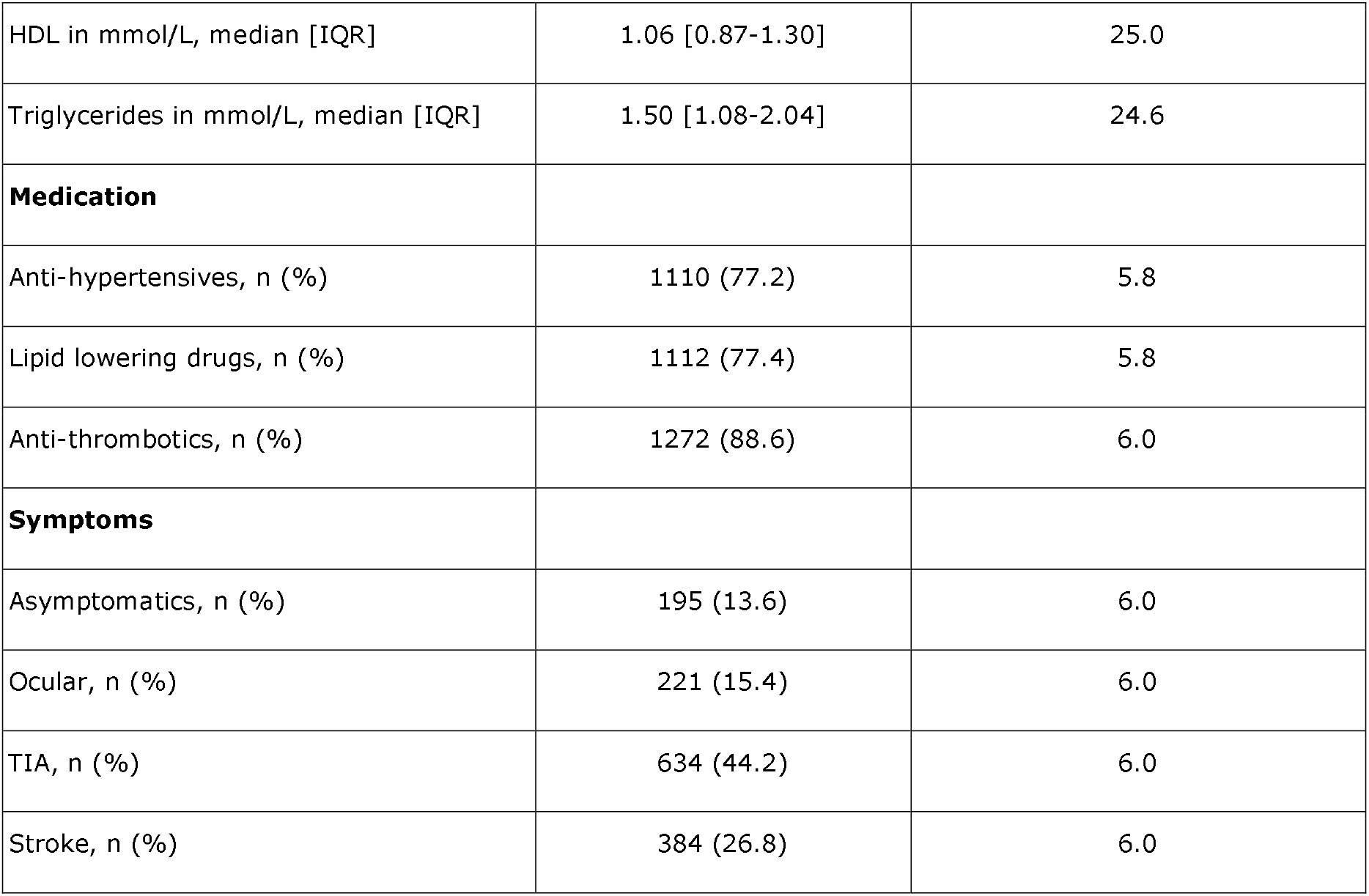
Baseline characteristics of genotyped CEA patients from the Athero-Express Biobank Study. Cerebrovascular disease history is defined by ischemic stroke prior to surgery. Coronary artery disease history includes diagnosed coronary artery disease, myocardial infarction, percutaneous coronary intervention, and coronary artery bypass grafting. Peripheral disease history includes diagnosed peripheral arterial occlusive disease, femoral artery interventions, and ankle-brachial index <70. Type 2 diabetes mellitus includes all individuals with diagnosed type 2 diabetes mellitus and those on appropriate medication. Hypertension includes all individuals with self-reported hypertension. Current smokers include all individuals smoking up to 6 months until the surgery date. BMI, kg/m^2^. eGFR rate was based on the Modification of Diet in Renal Disease formula, mL/min/1.73m^2^. Anti-hypertensives include all anti-hypertension medication. Anti-thrombotics include clopidogrel, dipyridamole, acenocoumarol, ascal, and other anti-platelet drugs. Missing data shows the percentage of the patients of which we lack information on the specific patient characteristic.

OSM is secreted by, among others, neutrophils(12), monocytes(11), macrophages(11) and T-cells(10), and acts through binding to OSMR and LIFR(14,42,43) in the arterial wall(7,44). Thus we queried data from the Genotype-Tissue Expression project (GTEx)(23) for SNPs that alter *OSM* expression in whole blood and *LIFR* and *OSMR* expression in arterial tissue. There were no significant eQTLs for *OSM*, but there were two eQTLs associated with altered *OSMR* (rs13168867) or *LIFR* (rs10491509) expression in arterial tissue. The C allele of rs13168867 is associated with decreased *OSMR* expression in the tibial artery (**Figure 1A**), and the A allele of rs10491509 is associated with increased *LIFR* expression in the aortic artery (**Figure 1B**). Cross-tissue meta-analysis showed that these variants have m-values >0.9 in both tibial and aortic artery tissue, indicating a high posterior probability that they are single *cis*-eQTLs in both tissues (**Supplemental Figure 1** and **2**).

**Figure 1:**
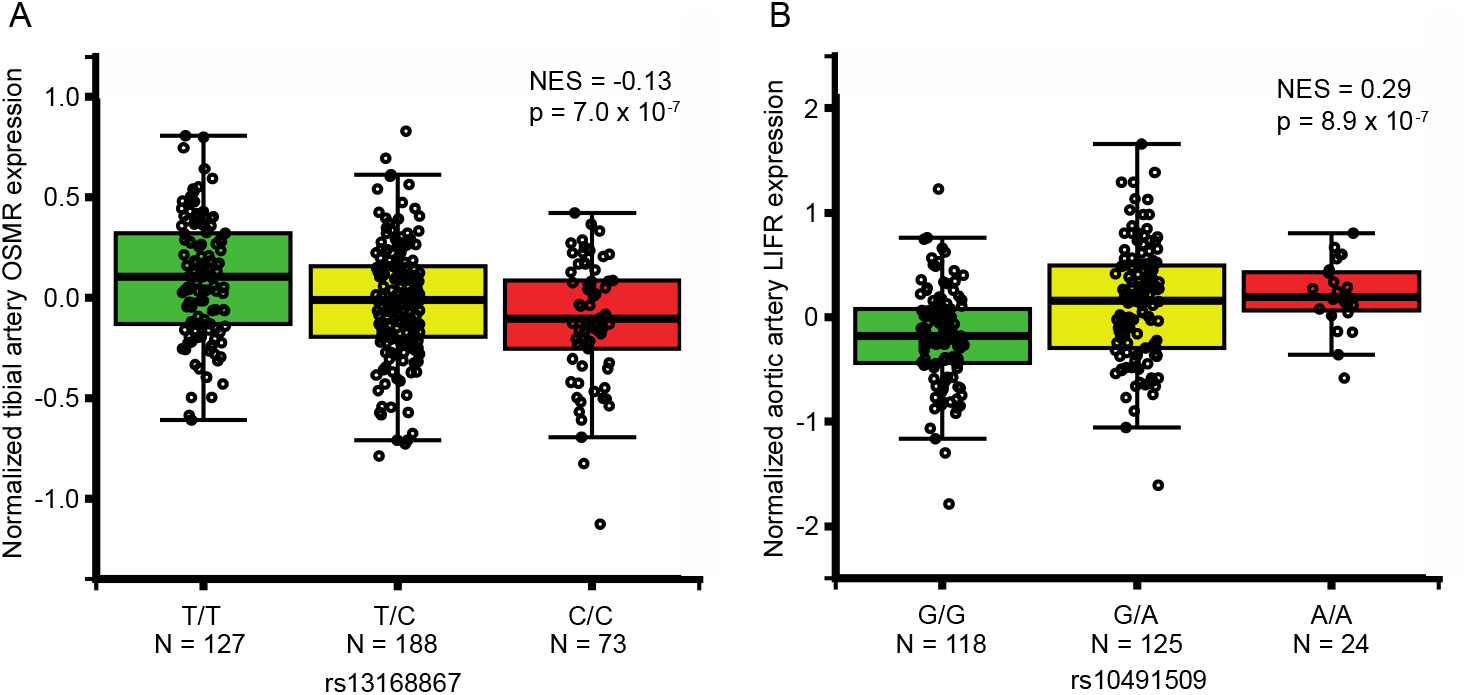
Association of OSMR and LIFR variants in non-diseased arterial tissues. Per variant, the normalized expression of OSMR **(A)** and LIFR **(B)** is given in non-diseased arterial tissue. Data from GTEx Portal (www.gtexportal.org). NES: Normalized effect size. In aortic arterial tissue, rs13168867 had a NES of -0.123 and in tibial arterial tissue, rs10491509 had a NES of 0.0881 (**Supplementary Figure 1** and **2**).

### 3.2 Genetic association with plaque vulnerability

To determine the effect of OSM signaling on the overall plaque vulnerability, we correlated the rs13168867 and rs10491509 genotypes to the overall plaque vulnerability, which was given a score ranging from 0 (least vulnerable plaque) to 4 (most vulnerable plaque). The effect allele of variant rs13168867 in the *OSMR* locus was significantly correlated with an increased overall plaque vulnerability (β=0.118 ± s.e.=0.040 (C allele), *p*=3.00×10^-3^, **Figure 2**). No association was observed with rs10491509 and overall plaque vulnerability.

**Figure 2:**
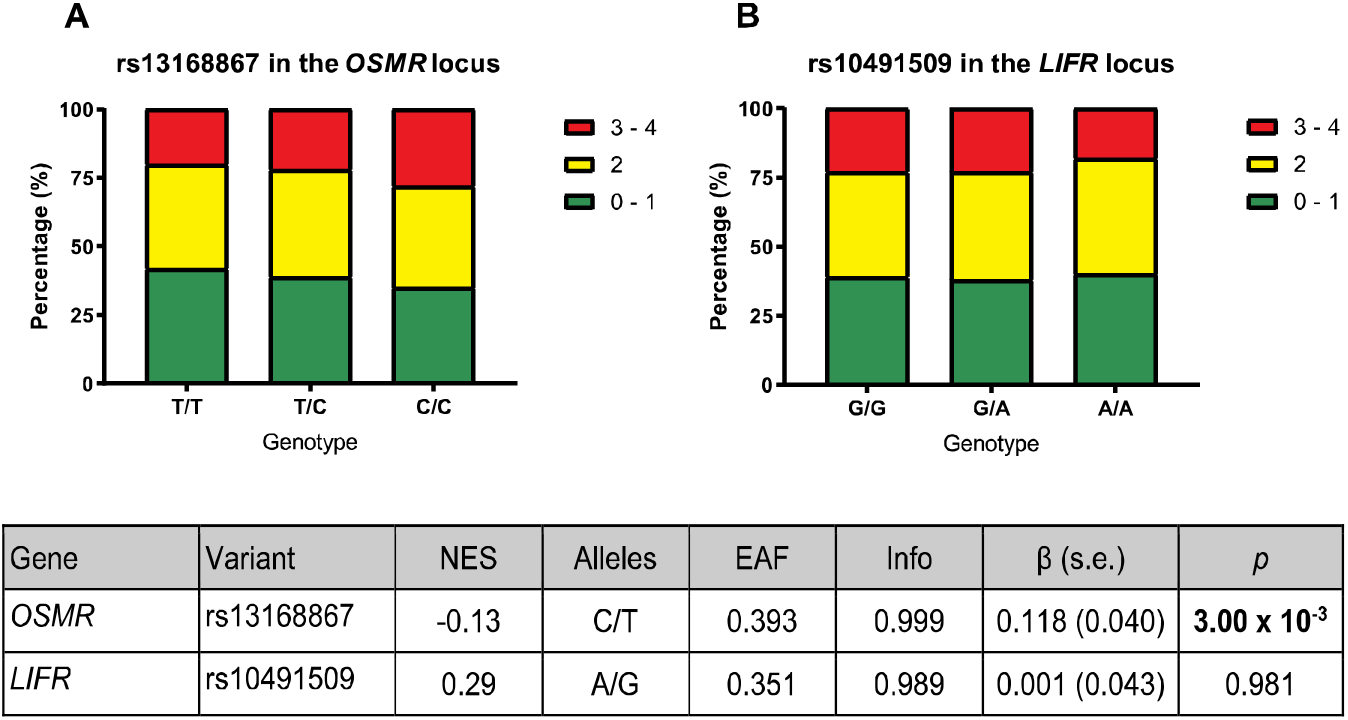
Association of OSMR and LIFR variants with overall plaque vulnerability. The plaques were given a vulnerability score ranging from 0 (least vulnerable) to 4 (most vulnerable). The bars represent the proportion of each plaque score per genotype for rs13168867 in the *OSMR* locus **(A)** and rs10491509 in the *LIFR* locus **(B)**. The table shows the results from GTEx Portal(23) where *Gene* is the gene of interest; *Variant* is the eQTL as found in GTEx Portal, and *NES* refers to the normalized effect size on expression from GTEx Portal. The *Alleles* refer to the effect allele and the other allele in both GTEx Portal and the Athero-Express Biobank. The remaining columns in the table are referring to the analysis of these eQTLs with respect to plaque vulnerability in the Athero-Express Biobank Study. Here *EAF* represents effect allele frequency; *Info* refers to the estimated imputation score. The effect size (β) and standard error (s.e.) are relative to the effect allele; *p* indicates the p-value of association with plaque vulnerability of the given eQTL; Bold *p*: p-value of association surpasses significance threshold (*p* <3.13×10^-3^).

### 3.3 Genetic association with plaque phenotypes

To determine the effect of OSM signaling on the individual plaque characteristics comprising the overall plaque vulnerability, we assessed the association between rs13168867 and rs10491509 and seven plaque phenotypes in the Athero-Express Biobank Study. Although not significant after correction for multiple testing, the strongest associations were observed between the effect allele of variant rs13168867 in the *OSMR* locus and intraplaque fat (β=0.248 ± s.e.=0.088 (C allele), *p*=4.66×10^-3^), and collagen content (β=-0.259 ± s.e.=0.095 (C allele), *p*=6.22×10^-3^, **Table 2**). No associations were observed between rs10491509 and any of the plaque phenotypes.

**Table 2:** *OSMR* and *LIFR* variants and their association with plaque phenotypes. For each variant, the association with plaque phenotypes is given. NES: the normalized effect size on expression; Alleles: the effect allele and the other allele, respectively; EAF: effect allele frequency in the Athero-Express Biobank; Info: estimated imputation score in the Athero-Express Biobank; β: effect size; s.e.: standard error; *p*: p-value of association; Adj. *p*: Bonferroni adjusted p-value. Calcification and collagen were classified as no/minor vs moderate/heavy, fat content as 10% vs >10% fat of plaque area, intraplaque hemorrhage was classified as absent vs present. Smooth muscle cells and macrophages were classified as Box-Cox transformed percentage of plaque area and vessel density as Box-Cox transformed number of vessels/hotspot. None of the p-values surpassed significance threshold (3.13×10^-3^).

| Gene | Variant | NES | Alleles | EAF | Info | Phenotype | $\beta$ (s.e.) | *p* |
| --- | --- | --- | --- | --- | --- | --- | --- | --- |
| *OSMR* | rs13168867 | -0.13 | C/T | 0.393 | 0.999 | Calcification | 0.036 (0.077) | 0.637 |
|  |  |  |  |  |  | Collagen | -0.259 (0.095) | 6.22 x 10<sup>-3</sup> |
|  |  |  |  |  |  | Fat content | 0.248 (0.088) | 4.66 x 10<sup>-3</sup> |
|  |  |  |  |  |  | Intraplaque hemorrhage | -0.014 (0.080) | 0.862 |
|  |  |  |  |  |  | Smooth muscle cells | 0.001 (0.011) | 0.913 |
|  |  |  |  |  |  | Vessel density | -1.06x10<sup>-4</sup> (0.004) | 0.976 |
|  |  |  |  |  |  | Macrophages | 0.004 (0.015) | 0.809 |
| LIFR | rs10491509 | 0.29 | A/G | 0.351 | 0.989 | Calcification | 0.046 (0.082) | 0.577 |
|  |  |  |  |  |  | Collagen | 0.134 (0.104) | 0.194 |
|  |  |  |  |  |  | Fat content | 0.086 (0.094) | 0.363 |
|  |  |  |  |  |  | Intraplaque hemorrhage | 0.071 (0.086) | 0.414 |
|  |  |  |  |  |  | Smooth muscle cells | -0.003 (0.012) | 0.840 |
|  |  |  |  |  |  | Vessel density | 0.002 (0.004) | 0.577 |
|  |  |  |  |  |  | Macrophages | 0.015 (0.016) | 0.354 |

### 3.4 Known eQTLs of OSMR and LIFR expression in non-diseased arterial tissue are not associated with expression in atherosclerotic plaques

Atherosclerotic disease progression changes the artery-specific transcriptional dynamics, and may therefore abolish the effects of known *OSMR* and *LIFR* eQTLs in non-atherosclerotic arterial tissues. Thus, we tested whether these eQTLs were associated with *OSMR* and *LIFR* expression in carotid atherosclerotic plaques. Neither variant showed associations with expression of *OSM, OSMR*, and *LIFR* (**Supplementary Table 1**).

### 3.5 Intraplaque *OSMR* expression is not associated with plaque vulnerability

As rs13168867 was associated with an increased overall plaque vulnerability, we next investigated if intraplaque *OSMR* expression levels associated with overall plaque vulnerability. Differential expression analyses, comparing the reference score (0) with each increasing vulnerability score (1, 2, 3, or 4) showed no associations between *OSMR* plaque expression levels and plaque vulnerability (**Supplemental Table 2**). Neither did intraplaque *OSM* or *LIFR* expression associate with plaque severity.

### 3.6 Known eQTLs of OSMR and LIFR do not associate with cardiovascular diseases

The Athero-Express comprises patients with advanced stage atherosclerotic plaques. Therefore, we assessed the effect of known *OSMR* and *LIFR* eQTLs on coronary calcification (CAC) as intermediate phenotype of atherosclerotic burden, and primary cardiovascular outcomes as clinical manifestation. We queried summary statistics from GWAS on CAC (n=2,674)(45), coronary artery disease (CAD, n=336,755)(45,46), and ischemic stroke subtypes (sample sizes 242,573-522,258)(45–47). Neither eQTL associated with increased CAC burden, or cardiovascular disease susceptibility (**Supplemental Table 3**).

## 4. Discussion

We investigated whether common variants associated to gene expression, i.e., eQTLs, near *OSM, OSMR* and *LIFR* affect overall plaque vulnerability and phenotype. We showed that one *cis*-acting eQTL (rs13168867, C/T), of which the C allele associates with reduced *OSMR* expression in non-diseased arterial tissue, significantly associates with increased plaque vulnerability after correction for multiple testing. This suggests that a decrease in *OSMR* expression and therefore possibly a decrease in OSM signaling, increases the chance of developing a vulnerable plaque.

To gain further insight into the role of genetically decreased *OSMR* expression on developing a vulnerable plaque, we examined the effect of rs13168867 on individual plaque characteristics in more detail. The strongest associations were found for rs13168867 with increased intraplaque fat and decreased collagen content, suggesting that reduced OSM signaling results in a larger lipid core and less fibrosis -in line with a more vulnerable plaque phenotype. We previously showed that OSM enhances intercellular adhesion molecule (ICAM)-1 expression on human endothelial cells(8). Reduced *OSMR* expression, which hypothetically results in reduced OSM signaling, may therefore result in reduced ICAM-1 expression. ICAM-1 depletion leads to M1 macrophage polarization(48), which is the pro-inflammatory macrophage subtype that promotes an unstable plaque phenotype(49). Reduced OSM signaling could also explain the decreased collagen content as OSM enhances in vitro fibroblast proliferation and collagen formation<u>(20)</u>. Moreover, it was previously shown that OSM enhances liver fibrosis in mice(50) and that OSM is upregulated in patients with pulmonary fibrosis(51). A reduction in OSM signaling caused by decreased OSMR expression may therefore result in decreased collagen content. Further studies are needed to investigate these hypotheses.

A possible explanation for the lack of associations for the variant (rs10491509) in the *LIFR* locus could be that an increase in *LIFR* expression would not affect OSM signaling as, hypothetically, there might already be a *LIFR* surplus and therefore, an increase in *LIFR* expression will not affect OSM signaling.

Although rs13168867 did associate with plaque vulnerability, no associations were found between rs13168867 and intraplaque *OSMR* expression, intraplaque *OSMR* expression and plaque vulnerability, nor did rs13168867 associate with cardiovascular disease outcomes. Possibly, OSM signaling mainly affects atherogenesis and atherosclerosis development in the initial phases of the disease. Arterial *OSMR* expression is reduced in human atherosclerotic plaques when compared to normal arteries^9^ and may therefore have bigger effects in the initial phase, when *OSMR* expression is still high. Another possible explanation is that OSM signaling may be overruled by for example, other cytokines in later stages of the disease. Lastly, although coronary thrombosis, and therefore cardiovascular disease, is most often caused by plaque rupture, which is most likely to happen in vulnerable plaques, thrombosis can also be triggered by other processes, including plaque erosion and atrial fibrillation(52,53). Xie *et al* showed that OSM is associated with thrombosis in patients with atrial fibrillation and suggested that OSM exerts thrombogenic effects by increasing tissue factor expression and decreasing the expression of tissue factor pathway inhibitors(53). So, OSM could potentially increase the risk of cardiovascular disease through its thrombogenic effects and at the same time decrease the risk of cardiovascular disease by its atheroprotective effects. Potentially, the seemingly atheroprotective effect of OSM that we described in our current study may be neutralized by the thrombogenic or potential other cardiovascular disease driving effects of OSM.

A limitation of association studies like ours, is that it is challenging to uncover the biological meaning of the discovered associations. It is likely that a reduction in OSMR expression on the arterial wall reduces OSM signaling, but this is difficult to verify. Firstly, OSM signaling is not only dependent on OSMR, but also on the blood OSM levels. If there is no or little OSM present in the blood, there might have been a surplus of OSMR and in this case, there will be no change in OSM signaling. Another possibility is that there is not only a decrease of OSMR expression on the arterial wall, but also a decrease in circulating OSMR levels, which can also bind to OSM and acts as a neutralizer(54), also resulting in no net difference in OSM signaling. Moreover, this study cannot make a distinction between the timing and the duration of OSM signaling, which may differentially affect atherosclerosis development as previous studies have shown that OSM, like IL-6, can act differently in the acute phase than in the chronic phase(8,9,55,56). Finally, we focused on only three genes (*OSM, OSMR* and *LIFR*), while atherosclerosis is a multifactorial disease. Although studies like ours can be very insightful to better understand the disease, single variants seldomly show big correlations with disease outcome.

Compared to genome-wide association studies that include thousands of individuals, the Athero-Express Biobank Study is relatively small (n=1,443), and, given its design, finite in size. However, it is well suited to examine the effect of common disease-associated genetic variation on plaque morphology and characteristics. Indeed, we estimated the power at ±75% given a MAF=0.40 (approximately the frequency of rs13168867) and relative risk=1.28 (http://csg.sph.umich.edu/abecasis/gas_power_calculator/).

Recent developments in single-cell expression analyses might extend on the present study by investigating which cell types, that are present in the plaque, most abundantly express *OSM, OSMR* and *LIFR*. Furthermore, it would be interesting to investigate if the *OSMR/LIFR* expression ratio correlates with plaque vulnerability and if this ratio might be a predictor of plaque vulnerability.

## 5. Conclusion

Based on this work we conclude that the variant rs13168867 in the *OSMR* locus is associated with increased plaque vulnerability, but not with coronary calcification or cardiovascular disease susceptibility. Given the multiple testing burden for individual plaque characteristics, it remains unclear through which precise biological mechanisms OSM signaling exerts its effects on plaque morphology, although our data point towards lipid metabolism and extracellular matrix remodeling. However, the OSM-OSMR/LIFR pathway is unlikely to be causally involved in lifetime cardiovascular disease susceptibility as none of the investigated eQTLs associated with cardiovascular diseases.

### Financial support

SWvdL is funded through grants from the Netherlands CardioVascular Research Initiative of the Netherlands Heart Foundation (CVON 2011/B019 and CVON 2017-20: Generating the best evidence-based pharmaceutical targets for atherosclerosis [GENIUS I&II]). This work was supported by ERA-CVD, grant number: 01KL1802. FWA is supported by UCL Hospitals NIHR Biomedical Research Centre. DvK, HP and DT are funded through the FP7 EU project CarTarDis (FP7/2007-2013) under grant agreement 602936. AB was funded through the Taxinomisis grant, part of the European Union’s Horizon 2020 research and innovation program (No 755320). HP received funding from the TNO research program “Preventive Health Technologies”. The funding sources were not involved in study design, collection, analysis and interpretation of data, writing of the report and in the decision to submit the article for publication.

## Author contributions

DvK: Conceptualization, Formal analysis, Writing -original draft. IvK: Data curation. AB: Conceptualization, Formal analysis, Writing – review & editing. HP: Writing – review & editing. AvG: Writing – review & editing. GJdB: Conceptualization. FA: Conceptualization. DT: Conceptualization, Writing – review & editing. GP: Conceptualization, Writing – review & editing. SvdL: Conceptualization, Formal analysis, Writing – original draft, review & editing.

## Acknowledgments

We would like to thank dr. Jessica van Setten and acknowledge her for imputing our datasets using an in-house developed imputation pipeline. Evelyn Velema and Petra Homoet-Van der Kraak are acknowledged for the immunohistochemical stainings. We also acknowledge the support from the Netherlands CardioVascular Research Initiative from the Dutch Heart Foundation, Dutch Federation of University Medical Centres, the Netherlands Organisation for Health Research and Development and the Royal Netherlands Academy of Sciences (“GENIUS I & II”, CVON2011-19) and the TNO research program “Preventive Health Technologies”.

