## Supplemental Material for "Common variants associated with *OSMR* expression contribute to carotid plaque vulnerability, but not to cardiovascular disease in humans"

Accompanying

by

D. van Keulen, MSc^1,2,3,4^, I. D. van Koeverden, MD PhD^1^, A. Boltjes PhD^2^, H. M. G. Princen, PhD^4^, A. J. van Gool, PhD^5,6^, G.J. de Borst, MD PhD^7^, F.W. Asselbergs, MD PhD^8,9,10^, D. Tempel, PhD^2,3,11^, G. Pasterkamp, MD PhD^2^*, S.W. van der Laan, PhD^2^*.

^1^Laboratory of Experimental Cardiology, University Medical Center Utrecht, University of Utrecht, Utrecht, The Netherlands; ^2^Laboratory of Clinical Chemistry and Hematology, University Medical Center Utrecht, University of Utrecht, Utrecht, The Netherlands; ^3^Quorics B.V., Rotterdam, The Netherlands; ^4^TNO-Metabolic Health Research, Gaubius Laboratory, Leiden, The Netherlands; ^5^Translational Metabolic Laboratory, Radboudumc, Nijmegen, The Netherlands; ^6^TNO- Microbiology & Systems Biology, Zeist, The Netherlands; ^7^Department of Vascular Surgery, University Medical Center Utrecht, University of Utrecht, Utrecht, The Netherlands; ^8^Department of Cardiology, Division Heart & Lungs, University Medical Center Utrecht, Utrecht University, Utrecht, The Netherlands; ^9^Institute of Cardiovascular Science, Faculty of Population Health Sciences, University College London, London, United Kingdom; ^10^Health Data Research UK and Institute of Health Informatics, University College London, London, United Kingdom; ^11^SkylineDx B.V., Rotterdam, The Netherlands.

* these authors contributed equally

### Supplemental Text

To be able to assess the global expression profile, plaque segments were thawed, cut up, and further homogenized using ceramic beads and tissue homogenizer (Precellys, Bertin instruments, Montigny-le-Bretonneux, France), in the presence of TriPure (Sigma Aldrich), and RNA was isolated according to TriPure manufacturer’s protocol. From here, RNA in the aqueous phase was precipitated using isopropanol, and washed with 75% ethanol, and subsequently stored in 75% ethanol for later use or used immediately after an additional washing step with 75% ethanol.

*Library preparation*

From here, library preparation was performed, adapting the CEL-Seq2 protocol for library preparation(34,35), as described before(36). After removing ethanol, and air-drying the pellet, primer mix containing 5ng primer per reaction was added, initiating primer annealing at 65°C for 5min. Subsequent RT reaction was performed; first strand reaction for 1h at 42°C, heat inactivated for 10m at 70°C, second strand reaction for 2h at 16°C, and then put-on ice until proceeding to sample pooling. The primer used for this initial reverse-transcription (RT) reaction was designed as follows: an anchored polyT, a unique 6bp barcode, a unique molecular identifier (UMI) of 6bp, the 5’ Illumina adapter and a T7 promoter, as described(36). Each sample now contained its own unique barcode, due to the primer used in the RNA amplification making it possible to pool together complementary DNA (cDNA) samples at 7 samples per pool. cDNA was cleaned using AMPure XP beads (Beckman Coulter), washed with 80% ethanol, and resuspended in water before proceeding to the *in vitro* transcription (IVT) reaction (AM1334; Thermo-Fisher) incubated at 37°C for 13 hours. Next, primers were removed by treating with Exo-SAP (Affymetrix, Thermo-Fisher) and amplified RNA (aRNA) was fragmented, and then cleaned with RNAClean XP (Beckman-Coulter), washed with 70% ethanol, air-dried, and resuspended in water. After removing the beads using a magnetic stand, RNA yield and quality in the suspension were checked by Bioanalyzer (Agilent).

cDNA library construction was then initiated by performing an RT reaction using SuperScript II reverse transcriptase (Invitrogen/Thermo-Fisher) according to the manufacturer's protocol, adding randomhexRT primer as random primer. Next, PCR amplification was done with Phusion High-Fidelity PCR Master Mix with HF buffer (NEB, MA, USA) and a unique indexed RNA PCR primer (Illumina) per reaction, for a total of 11-15 cycles, depending on aRNA concentration, with 30 seconds elongation time. PCR products were cleaned twice with AMPure XP beads (Beckman Coulter). Library cDNA yield and quality were checked by Qubit fluorometric quantification (Thermo-Fisher) and Bioanalyzer (Agilent), respectively. Libraries were sequenced on the Illumina Nextseq500 platform; paired end, 2 x 75bp.

*Mapping*

Upon sequencing, retrieved fastq files were de-barcoded, split into forward and reverse reads. Subsequently, these were mapped making use of Burrows-Wheel aligner (BWA(37)) version 0.7.17-r1188, calling ‘bwa aln’ with settings -B 6 -q 0 -n 0.00 -k 2 -l 200 -t 6 for R1 and -B 0 -q 0 -n 0.04 -k 2 -l 200 -t 6 for R2, ‘bwa sampe’ with settings -n 100 -N 100, and a cDNA reference (assembly hg19, Ensembl release 84). Read counts and UMI counts were derived from SAM files using custom perl code, and then gathered into count matrices. Genes were annotated with Ensembl ID’s, and basic quality control was performed, encompassing filtering out samples with low gene numbers (<10,000 genes), and read numbers (<18,000 reads). These steps resulted in 641 samples with up to 60,674 genes (Ensembl ID’s), and median of 178,626 reads per sample.

### Supplemental Figures

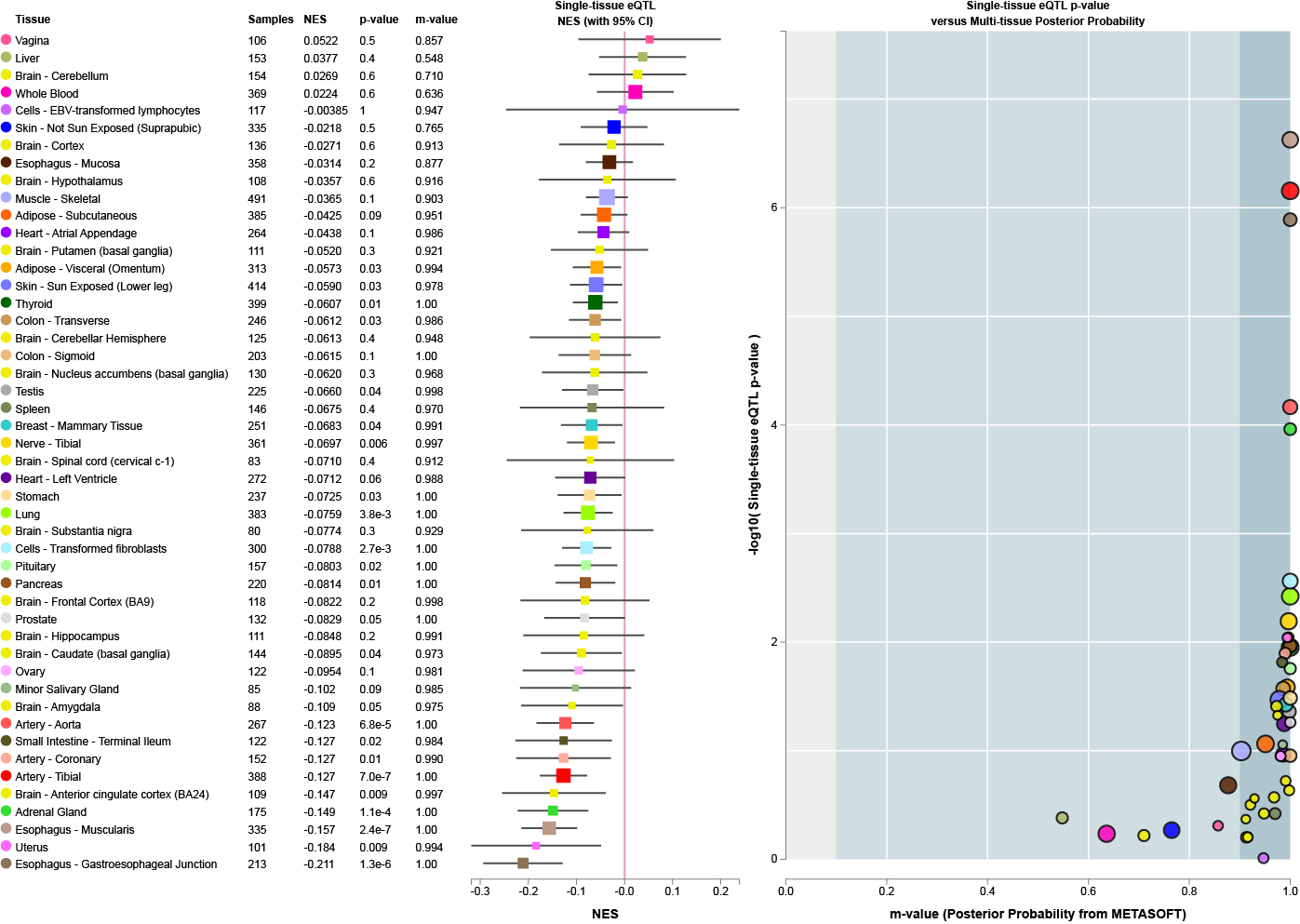

**Supplemental Figure 1. eQTL analysis of rs13168867 with *OSMR* expression in multiple tissues from the GTEx Project.** The forest plot shows the correlation of rs13168867 with *OSMR* expression per tissue and the posterior probability (m-value) from METASOFT^1^ showing the posterior probability that the specific eQTL exists in each tissue (a large m-value indicates that the variant is predicted to be an *cis*-acting eQTL for *OSMR* in that specific tissue). Data is obtained from GTEx Portal^2,3^.

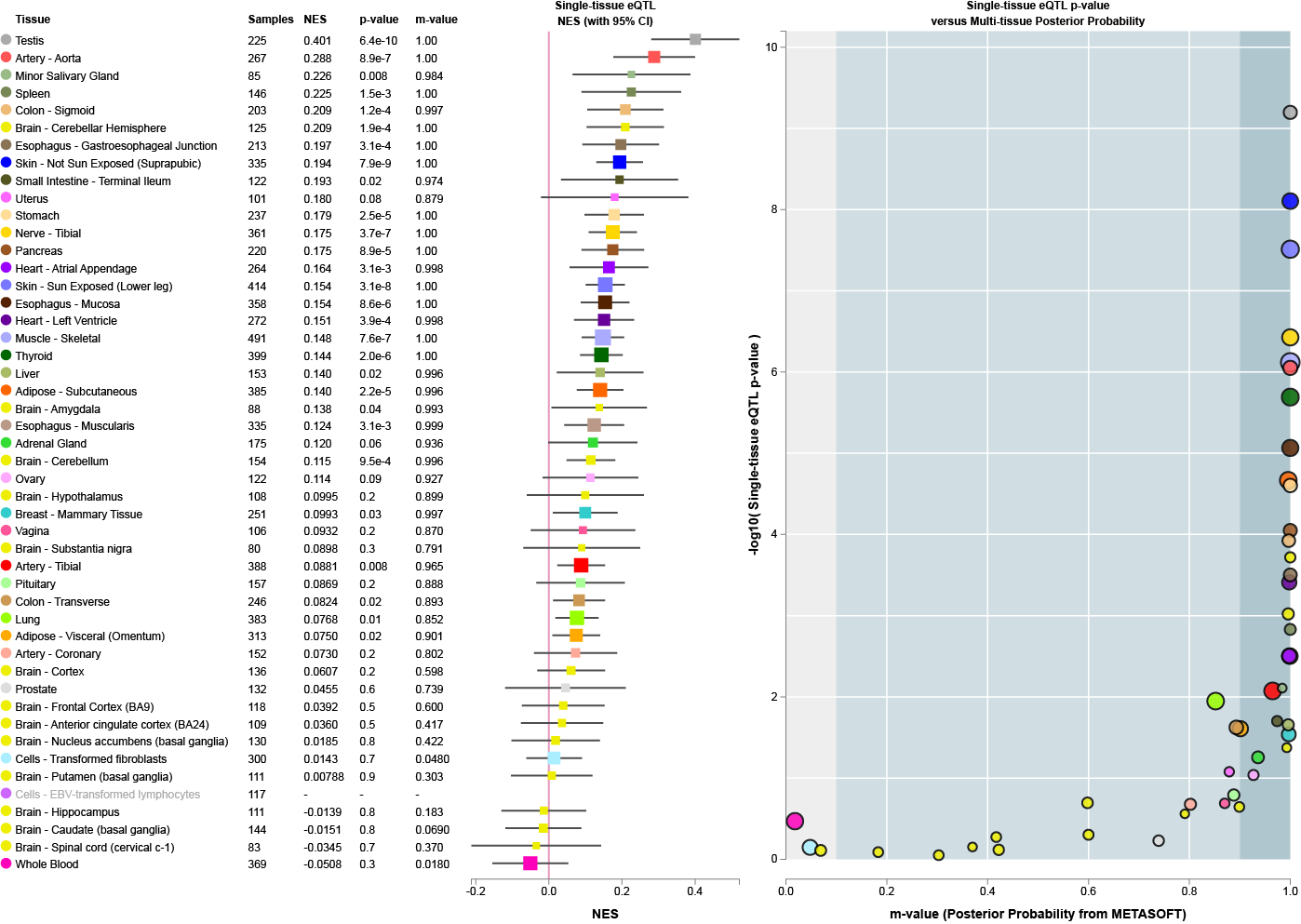

**Supplemental Figure 2. eQTL analysis of rs10491509** **with *LIFR* expression in multiple tissues from the GTEx Project.** The forest plot shows the correlation of rs10491509 with *LIFR* expression per tissue and the posterior probability (m-value) from METASOFT^1^ indicating that the variant is predicted to be an *cis*-acting eQTL for *LIFR* in that specific tissue. Data is obtained from GTEx Portal^2,3^.

### Supplemental Tables

**Supplementary Table 1: *OSMR* and *LIFR* variants and their association with intraplaque *OSM, OSMR* and *LIFR* expression.** For each Gene affected by an eQTL in arterial tissue from GTEx, the association with intraplaque *OSM, OSMR* and *LIFR* expression is given. Alleles: the effect allele and the other allele, respectively; β: effect size; s.e.: standard error; *p*: p-value of association.

| **Gene** | **eQTL** | **Alleles** | **Gene in plaque** | **β (s.e.)** | ***p*** |
| --- | --- | --- | --- | --- | --- |
| *OSMR* | rs13168867 | C/T | *OSM* | -0.035 (0.097) | 0.724 |
|  |  |  | *OSMR* | -0.002 (0.033) | 0.961 |
|  |  |  | *LIFR* | -0.015 (0.034) | 0.648 |
| *LIFR* | rs10491509 | A/G | *OSM* | -0.103 (0.103) | 0.323 |
|  |  |  | *OSMR* | 0.019 (0.035) | 0.589 |
|  |  |  | *LIFR* | 0.014 (0.035) | 0.692 |

**Supplemental Table 2: Differential expression analysis of the reference overall plaque vulnerability score (0) vs. others (1, 2, 3, or 4).** *Gene*: HGNC gene symbol. *EnsemblID*: Ensembl 84 gene ID. *baseMean*: mean gene expression, i.e. normalized count divided by size factors, over all samples. *log_2_FC*: effect size on a log_2_-scale. *SE*: standard error of the effect size. *p*: p-value

|  | **Gene** | **EnsemblID** | **baseMean** | **log_2_FC** | **SE** | ***p*** |
| --- | --- | --- | --- | --- | --- | --- |
| **results 0vs1** | *OSMR* | ENSG00000145623 | 4.091 | 0.207 | 0.123 | 0.091 |
|  | *LIFR* | ENSG00000113594 | 2.442 | 0.138 | 0.151 | 0.340 |
|  | *OSM* | ENSG00000099985 | 0.085 | -0.025 | 0.142 | 0.966 |
| **results 0vs2** | *OSMR* | ENSG00000145623 | 4.091 | 0.128 | 0.113 | 0.237 |
|  | *LIFR* | ENSG00000113594 | 2.442 | 0.017 | 0.139 | 0.783 |
|  | *OSM* | ENSG00000099985 | 0.085 | 0.023 | 0.152 | 0.845 |
| **results 0vs3** | *OSMR* | ENSG00000145623 | 4.091 | 0.132 | 0.126 | 0.280 |
|  | *LIFR* | ENSG00000113594 | 2.442 | 0.125 | 0.155 | 0.397 |
|  | *OSM* | ENSG00000099985 | 0.085 | 0.043 | 0.132 | 0.731 |
| **results 0vs4** | *OSMR* | ENSG00000145623 | 4.091 | 0.182 | 0.141 | 0.197 |
|  | *LIFR* | ENSG00000113594 | 2.442 | 0.334 | 0.171 | 0.050 |
|  | *OSM* | ENSG00000099985 | 0.085 | 0.029 | 0.113 | 0.771 |

**Supplemental Table 3: Association of OSMR and LIFR variants with cardiovascular outcomes and carotid IMT.**

*Phenotype*: *CAD*, coronary artery disease^4^; *CAC*, coronary artery calcification^5^; *AS*, all cause ischemic stroke^6^ IS*,* ischemic stroke^6^; *LAS*, large artery ischemic stroke^6^; *CES*, cardio-embolic stroke^6^; *SVD*, small vessel disease^6^. *eQTL*: variant of interest affecting expression of *OSMR* (rs13168867) and *LIFR* (rs10491509). *Position*: the chromosomal base pair position. *Alleles*: effect allele associated with trait susceptibility, and the other allele. *EAF*: effect allele frequency. *OR* (95% CI), odds ratio of association with 95% confidence interval. *p*: p-value of association. *N*: total sample size. *N cases*: number of cases. *N ctrls*: number of controls. * These are the *maximum* numbers of cases and controls as mentioned in Supplementary Figure 2 in the manuscript by Nelson *et al*.^4^. Thus, these numbers represent more samples in total than the column *N* reports in the summary statistics as downloaded from the website (http://www.cardiogramplusc4d.org/data-downloads/). This has to do with per-cohort missing data for these particular variants.

| **Phenotype** | **eQTL** | **Position** | **Alleles** | **EAF** | **OR (95%CI)** | ***p*** | **N** | **N cases** | **N ctrls** |
| --- | --- | --- | --- | --- | --- | --- | --- | --- | --- |
| CAD | rs13168867 | chr5:38,975,458 | T/C | 0.60 | 1.01 (1.00-1.03) | 0.17 | 336,755 | 113,937* | 339,115* |
| CAC |  |  |  | 0.42 | 1.11 (0.96-1.29) | 0.16 | 2,674 | NA | NA |
| AS |  |  |  | 0.58 | 1.00 (0.98-1.01) | 0.60 | 522,258 | 67,030 | 455,228 |
| IS |  |  |  | 0.58 | 1.00 (0.98-1.01) | 0.83 | 511,594 | 60,341 | 451,253 |
| LAS |  |  |  | 0.57 | 0.96 (0.93-1.00) | 0.07 | 245,201 | 6,688 | 238,513 |
| CES |  |  |  | 0.58 | 1.01 (0.98-1.04) | 0.62 | 361,858 | 9,006 | 352,852 |
| SVD |  |  |  | 0.56 | 1.00 (0.97-1.03) | 0.98 | 298,777 | 11,710 | 287,067 |
| CAD | rs10491509 | chr5:38,472,617 | A/G | 0.33 | 1.00 (0.99-1.02) | 0.62 | 334,545 | 113,937* | 339,115* |
| CAC |  |  |  | 0.32 | 1.03 (0.88-1.20) | 0.76 | 2,674 | NA | NA |
| AS |  |  |  | 0.34 | 1.01 (1.00-1.03) | 0.11 | 519,435 | 64,821 | 454,614 |
| IS |  |  |  | 0.34 | 1.01 (0.99-1.03) | 0.21 | 507,648 | 58,546 | 449,102 |
| LAS |  |  |  | 0.35 | 1.03 (0.99-1.08) | 0.16 | 242,573 | 6,370 | 236,203 |
| CES |  |  |  | 0.34 | 1.00 (0.96-1.04) | 1.00 | 357,275 | 8,605 | 348,670 |
| SVD |  |  |  | 0.36 | 0.99 (0.96-1.03) | 0.64 | 296,151 | 11,203 | 284,948 |
